# Evolution of divergent daily temporal niches shaped by male-male competition can generate sympatric speciation

**DOI:** 10.1101/2024.07.31.601896

**Authors:** Titouan Bouinier, Arthur Brunaud, Charline Smadi, Violaine Llaurens

**Affiliations:** Centre Interdisciplinaire de Recherche en Biologie (CIRB), Collège de France, INSERM, CNRS, 11 Place Marcellin Berthelot, 75005 Paris, France; Institut de Systématique, Evolution, Biodiversité (ISYEB), Muséum national d’Histoire naturelle, CNRS, Sorbonne Université, EPHE, Université des Antilles CP 50, 57 rue Cuvier, 75005 Paris, France; LESSEM, INRAE, Univ. Grenoble Alpes, F-38402 St-Martin-d’Hères, France; Institut Fourier, CNRS, Univ. Grenoble Alpes, 38610 Gières, France

**Keywords:** Genetic incompatibility, Population divergence, Competition, Sex-ratio, Diel activity

## Abstract

Specialisation into different ecological niches is assumed as an important driver of speciation in sympatry. Here we focused on male-male competition fuelling population divergence, without assuming any ecological specialisation. We investigated how antagonistic interactions between males can promote divergent evolution of timings of reproductive activities during the day, as observed in some closely-related insect species living in sympatry. We used a multi-locus, comprehensive stochastic model to investigate the evolution of (1) the timing of reproductive activity as a quantitative trait and (2) neutral loci that may generate genetic incompatibilities among divergent individuals. We specifically explore how male-male competition for female access can generate negative frequency-dependence on the timing of reproductive activities and fuel population divergence. Our simulations in finite populations highlight the strong effect of male-male competition and operational sex-ratio on the evolution of divergent temporal niches. They also show how genetic incompatibilities promote the differentiation among populations with divergent temporal niches, but may also impair their co-existence. Our model therefore highlights male-male competition as an important factor shaping the evolution of diel niches, that may fuel sympatric speciation.

## Introduction

The coexistence of multiple species in sympatry is usually thought to be linked to their partition into distinct ecological niches (J. M. Smith, 1966). Within communities, species can be partitioned into different temporal niches, with variations in breeding season (*e.g.* in birds, Karr, 1976) or timing of daily activities (*e.g.* in tropical beetles, De Oliveira Ribeiro et al., 2022 or Lepidoptera, Beck and Linsenmair, 2006). Plants flowering time is a common example of differentiated reproductive timings, fuelling genetic differentiation between closely-related, sympatric species, as for instance in *Howea* palms (Savolainen et al., 2006), or *Gymnadenia* orchids (Soliva and Widmer, 1999). Temporal niches can be shaped by abiotic factors such as temperature, but also by interactions with closely-related species (see Taylor and Friesen, 2017 for a review). In particular, reproductive interference between closely related species might favour the evolution of divergent phenology in sympatry (Devries et al., 2008). In turn, temporal segregation would also contribute to the speciation process itself, because of preferential reproduction among individuals sexually active at the same period of time (Devaux and Lande, 2009). Allochronic speciation is often assumed to stem from the coupling of the activity timing trait with another ecological trait (Tessnow et al., 2022; Van Doorn et al., 2024). Nevertheless, multiple cases of potential allochronic speciation have been reported where no discernable ecological specialisation is observed between temporally differentiated species (Levitan et al., 2004; Pike et al., 2003). The conditions facilitating such allochronic speciation (Hendry and Day, 2005) in the absence of other ecological specialisation are still mostly unexplored, despite the repeated observation of temporal partitioning of species activities throughout the day, as for instance in tropical insects (see Caveney et al., 1995; De Oliveira Ribeiro et al., 2022; Devries et al., 2008; Lamarre et al., 2015).

Male-male competition for accessing available females is probably a key factor shaping the evolution of divergence in males timing of activities. For instance, the evolution of protandry promoted by selection linked to limited female availability has indeed been documented in a number of plant (Forrest, 2014; Zhao et al., 2020) and animal species (Morbey and Ydenberg, 2001). In species with separated sexes, male-male competition can indeed be strong, especially when the adult sex-ratio is unbalanced or when there is an asymmetry in the frequency of mating between males and females (Arak, 1983; Sullivan-Beckers and Cocroft, 2010). Furthermore, when female availability is a limiting factor for male reproduction, intense competitive interactions may occur among males and the evolution of non-overlapping timing of courtship may then reduce the deleterious effects of male-male competition. Nonetheless, whether temporal segregation could be an evolutionary response to male-male competition is still not clear (Ekrem et al., 2025). Competitive interactions among males may generate negative frequency dependent selection on the timing of reproductive activities (Doebeli, 1996). Here, we specifically aimed at investigating the effect of male-male competition on the evolution of temporal niches in absence of ecological divergence between these niches. We thus tested in which conditions the temporal divergence acting as pre-zygotic barrier may promote divergence in sympatry, without any ecological coupling.

Nonetheless, post zygotic barriers, such as hybrid viability, might also play an important role in allochronic speciation processes. Notably, disentangling whether temporal segregation is the result of a reinforcement process caused by the poor hybrid fitness in cases of secondary contact between species, or whether such segregation is the initial driver of speciation is an important question. Allochrony was shown to be an important factor fuelling population divergence, even when the initial trigger of divergence stems from another ecological factor (*e.g.* host shift, Tabuchi and Amano, 2003), suggesting an effect of allochrony in the reinforcement process.

Pre-zygotic isolation caused by allochronic migrations has also been suggested to reinforce genetic divergence between populations of the butterfly *Danaus chrysippus* (D. A. S. Smith et al., 2010). The level of post-zygotic isolation between sympatric species with divergent temporal niches is frequently unknown, but the accumulation of Bateson-Dobzhansky-Muller (*BDM* hereafter) incompatibilities (Orr and Turelli, 2001) is likely to interact with the preferential reproduction between synchronously active individuals, therefore facilitating the emergence of reproductive barriers between asynchronous populations. Such interactions between pre and post-zygotic barriers have been shown to fuel speciation by reinforcement in a model of neutral speciation triggered by isolation by distance (Baptestini et al., 2013). How much the interaction between genomic incompatibilities and temporal isolation fuels speciation is still an open question.

Here, we built a general model to investigate the effect of male-male competition on the evolution of the temporal window of daily reproductive activities and its impact on the population divergence and speciation. We assumed a single haploid species, with two separate sexes, obligatory sexual reproduction and asymmetry in the mating limitations between males and females (limited female reproduction *vs.* unlimited male reproduction). We modelled a quantitative trait of timing of sexual activity controlled by a multilocus architecture, in a uniform environment without any ecological selection. We also investigated the interaction between the evolution of this timing of activity and genetic incompatibilities by simulating the evolution of biallelic haploid loci within the genome. These loci either (1) had no impact on reproductive success or (2) prevented reproduction between genetically divergent individuals as they accumulated incompatibilities. We therefore estimated the contributions of pre- and/or post-zygotic factors in driving the emergence of divergent populations that differ in their reproductive timing, in the absence of any ecological specialisation.

## Methods

### Model overview

We developed a stochastic, individual-based birth–death model to study the evolution of daily reproductive activity timing and its consequences for allochronic divergence, assuming different levels of post-zygotic genetic incompatibilities. Simulations track the fate of individuals over a long sequence of identical days, without explicit seasonal or annual structure.

Each individual is characterized by a set of traits (Table 1) that determine its survival and reproductive success. In particular, both males and females express a heritable activity-timing trait *h_a_* ∈ [0, 1], which defines the daily timing of their reproductive activity. The trait *h_a_*is controlled by *B* haploid, unlinked loci with additive effects of equal magnitude 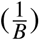, such that mutations at a single locus change *h_a_* by 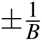. Free recombination among loci is assumed, and mutations occur at rate *µ* per locus.

**Table 1:** Individual attributes.

| Attribute | Description | Values |
| --- | --- | --- |
| *sex* | Sex of the individual | "M" or "F" |
| *h<sub>a</sub>* | Center of the reproductive activity period | List of *B* loci (with values 0/1) |
| *w<sub>a</sub>* | Range of the reproductive activity period | 0.15 |
| *g<sub>a</sub>* | Incompatibility loci | List of *B* loci (with values 0/1) |
| *E* | Number of eggs laid by a mated female | Variable positive integer |

The model, summarized in fig. 1, was inspired by the life cycle of Lepidoptera species, where immature larvae stages and sexually mature adult stage (imago) are strictly separated by the pupal stage. It can nevertheless be generalized to hemimetabolous and holometabolous insects (insects with multiple stages of development), as they generally display life cycles with stereotypical timing of transitions between developmental stages. In the model, immature stages are not represented explicitly; instead, density-dependent survival is applied to larvae, and these surviving offspring emerge as adults later the same day.

**Figure 1:**
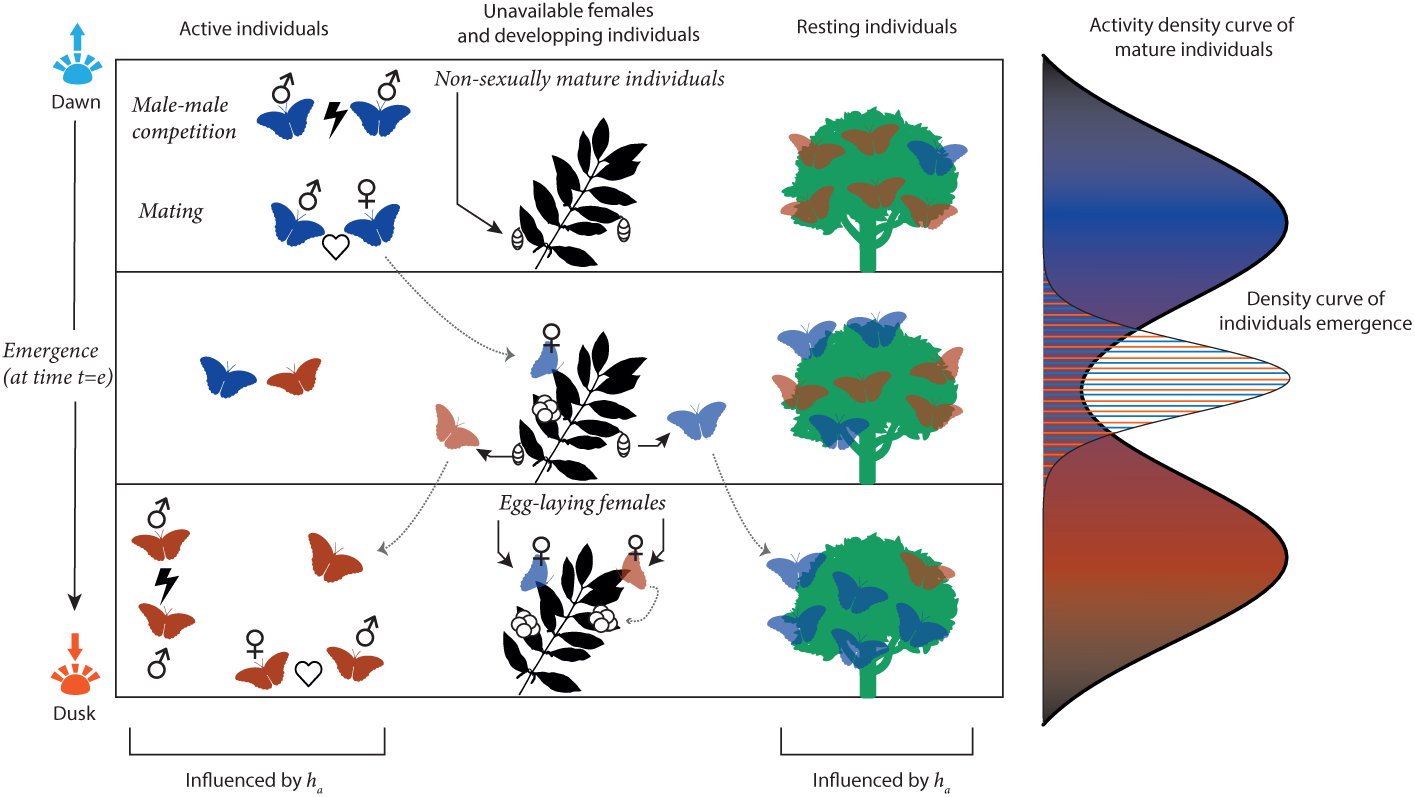
Schematic view of the model describing variations in activity patterns in a butterfly population, resulting in the evolution of two sub-populations, with individuals with early and late adult reproductive activity times represented by blue and orange butterflies respectively. The scheme describes a typical day in our simulation, where two sub-populations have successfully differentiated. Males active at the same time can participate to male-male competition, and mate with females. Once mated *p* times, females become unreceptive for reproduction and lay eggs that will give rise to new individuals. These non-sexually mature offspring (represented as pupae on the scheme) will become mature adults during the same day. During the period determined by the emergence time *t = e* each day, sexually mature adults can emerge from the previously produced pupae. Note that if an individual activity timing is later than the emergence timing, it will be become active within the same day, potentially mating right after emergence. In contrast, individuals emerging after their activity timings will start mating only the following day.

Time is structured into discrete days, each represented as a continuous interval from dawn (*t* = 0) to dusk (*t* = 1). Adult individuals are active only during a daily activity window of fixed width 2 ∗ *w_a_*, centered on their individual value of *h_a_*. Individuals can mate only when both partners are active simultaneously. Adult emergence occurs once per individual at a time drawn from a truncated normal distribution with mean *e* and variance *v_e_*, bounded to the interval [0, 1].

The simulation process used here follows the Doob-Gillespie method (see Gillespie, 1977; Tran, 2006), that allows to simplify the way individual behaviors actually happen during the course of our stochastic simulations, based on their respective probabilities. The probabilities of different events to happen depends on the individual attributes (Table 1), the parameters of the model (Table 2) and the population state. We describe below the sequence of events that may occur within and between days (further details regarding the simulation process are provided in the Supplementary Materials and Methods).

**Table 2:** Parameters of the model and baseline values used in stochastic simulations.

| Parameter | Description | Values |
| --- | --- | --- |
| $\beta$ | Encountering rate between active males and available females | **3** |
| $d$ | Basal death rate | **0.1** |
| $\delta_e$ | Mate searching energetic cost for all individuals | **0.1** |
| $\delta_c$ | Competition cost in active males | 0, 0.05, **0.1**, ..., 0.025 |
| $\delta_l$ | Egg laying cost in mated females | **0.1** |
| $p$ | Maximum number of partners per female | 1, 2, 3, 4, 5 |
| $b$ | Mean offspring number per female per day | **5** |
| $e$ | Emergence peak time | 0.1, 0.2, ..., **0.5**, ..., 0.9 |
| $v_e$ | Variance in the timing of emergence | **0.05** |
| $\mu$ | Mutation rate of loci | **0.03** |
| $B$ | Number of loci for each trait (genetic incompatibility, activity timing) | **50** |
| $G$ | Genetic threshold of reproductive isolation | 0.1, 0.2, ..., 0.8, 0.9, **1** |
| $K_{larvae}$ | Carrying capacity of the environment for immature offsprings (*e.g.* larvae) | **5000** |

### Daily schedule of events

Each simulation consists of a sequence of days indexed from 1 to 500. Within each day, time is continuous (*t* ∈ [0, 1]), with a succession of multiple random events (death, mating, emergence), while some processes occur synchronously at day boundaries (offspring/egg production).

#### 1. Dawn (*t* = 0): offspring production

All mated females that are alive at the start the day lay eggs. In cases where the mother has mated with multiple males (in simulations with *p >* 1), a sire will be chosen uniformly at random to fertilize the eggs. Density-dependent survival is applied to these eggs, producing a number of surviving offspring per female. Surviving offspring immediately enter the pupal pool and emerge as adults during the same day. The number of surviving offspring produced by each female is drawn from a Poisson distribution with parameter:

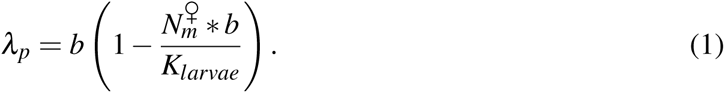

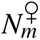 is the number of mated females, with *b* the number of eggs per female. *K_larvae_* is the carrying capacity of the immature stage. Density-dependent competition thus acts exclusively during the immature phase. Females lay eggs every day after their first mating, until death.

#### 2. Adult emergence (*t* ∼ *N*(*e*, *ve*))

Surviving offspring emerge as adults later the same day, at a time drawn from a truncated normal distribution *N*(*e*, *v_e_*) bounded to [0, 1]. We considered a balanced sex-ratio between males and females at emergence.

Newly emerged adults become sexually active immediately if their activity window overlaps the remaining portion of the day. Otherwise, they can only engage in reproductive activity starting the following day.

#### 3. Within-day dynamics (0 *< t <* 1)

During the day, adult individuals experience continuous-time stochastic events, whose rates are continuously computed:

- **Mating:** Active males and active receptive females mate at a rate proportional to their numbers and an encounter rate *β* (e.g. encounters between an active male and an active receptive female happens at a rate of *β* ∗ *N_active males_* ∗ *N_active_*_,*receptive*_ *_f emales_*). Males can mate multiple times, while females can mate up to *p* times, after which they become permanently unreceptive. Mating events occurring during day *d_n_* affect the paternity of offspring laid at dawn of day *d_n_*_+1_.
- **Mortality:** All adults experience a baseline death rate *d*. Additional mortality costs apply during periods of activity with an activity-related cost for both sexes, proportional to the amount of activity timing for each individuals (at a rate *δ_e_*). We also implemented sex-specific death rates: a behavioural competition cost for males, dependent on the current number of simultaneously active males (at a rate *δ_c_*), and a reproduction-dependent cost for females, which is proportional to the number of eggs laid (at a rate *δ_l_*).

#### 4. End of day (*t* = 1): carry-over

Individuals that survive until dusk persist into the next day, where they may be active again, along with newly emerged individuals at *t* ∼ *N*(*e*, *ve*).

### Offspring viability and genetic incompatibilities

To model post-zygotic reproductive barriers, individuals carry an additional set of *B* haploid, biallelic loci that may affect offspring viability but do not influence adult survival or reproductive performance. These loci are assumed to be unlinked and ecologically neutral, and are hereafter referred to as genetic incompatibility loci.

The viability of offspring produced by a given parental pair depends on the genetic distance between their genotypes at these loci. Genetic distance is quantified using the normalized Hamming distance:

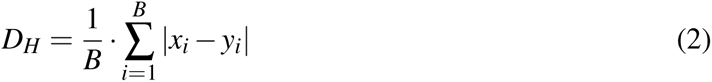

with *x_i_* and *y_i_* the allele values at the locus i for the individuals *x* and *y* respectively.

We assume a threshold-based incompatibility model, such that offspring produced by parental pairs with *D_H_ > G* are inviable. Depending on the value of *G*, we can simulate an absence (for *G* = 1) or varying degrees of post-zygotic barriers (for 0 ≤ *G <* 1). Although this formulation represents a simplified description of post-zygotic incompatibilities, it provides a tractable way to implement reproductive isolation between genetically divergent individuals and has been commonly used in previous theoretical studies (de Aguiar et al., 2009; Yamaguchi and Iwasa, 2017).

#### Inheritance and mutation

For viable offspring, alleles at both activity-timing loci and genetic incompatibility loci are inherited under free recombination. Each locus mutates independently with probability *µ*, with mutations resulting in a switch of allelic state (from 0 to 1 or 1 to 0).

### Stochastic simulations and equilibria

We ran each simulation for *T* = 500 days for each repetition, ensuring that the equilibrium is reached for simulations, and used a range of reasonable parameters to ensure the emergence and coexistence of differentiated sub-populations. Parameters and attribute range values are provided in Tables 1 and 2. The operational sex-ratio ranged from an average of 0.714 ± 0.08 to 0.36 ± 0.05 over the range of strictly positive *δ_c_* values tested in our model (assuming *G* = 1, *p* = 1). Similarly, mean lifetime of males ranged from an average of 0.71 ± 0.01 to 0.31 ± 0.06 day, and mean lifetime of females ranged from an average of 4.14 ± 0.01 to 4.12 ± 0.01 days. All simulations ended after 500 days, or when the entire population became extinct. The initial conditions for all simulations are a population of one hundred of individuals with a balanced sex-ratio, where all females are initially virgin. The initial genotypes at the loci controlling *h_a_* of all individuals are set to a identical assortment of 0 and 1, such that their initial phenotypic value is 0.5. The genetic incompatibility loci are initialized with values all equal to 0. A hundred replicates were run for each condition, with the range of parameters used is given in Table 2. Python (3.11.7) was used for modeling purposes and data cleaning, using the packages ”random”, ”numpy”, ”scipy” and “seaborn”. R (4.4.0) was used to plot all results, using the “ggplot2” package.

### Summary statistics drawn from simulation outcomes

To detect whether differentiated populations evolved during the course of a simulation, we tracked the distribution of the trait *h_a_*in the simulated populations and aim at detecting the number of peaks in this distribution. In Python (3.11.7), we used the *find_peaks* function from the “scipy.signal” package (Virtanen et al., 2020), to detect peaks and identify their position to define sub-populations. We used the minimum between the two peaks in the reproductive activity distribution curves to determine to which sub-population each individual belonged to. Individuals with *h_a_* values greater or lesser than this minima are sorted to either sub-population. Note that this minimum is not necessarily close to 0, meaning that some overlap can still occur between sub-populations. A group of individuals with similar timing of activities that diverge from the ancestral population is then considered as an actual differentiated sub-population only when (1) it lasted more than forty days within the simulation and when (2) the position of the peak within the differentiated sub-population did not vary by more than 0.15 units of time over a day. Those conservative requirements were determined from previous simulations, when searching for parameters that would guarantee large enough sub-population sizes and stable sub-populations over the length of a simulation. We must note that in our simulations, we only observed cases with either one or two peaks, (*i.e.* either one or two differentiated sub-populations), no matter the set of parameters used.

We considered three categories of sub-populations in our model:

1. *Immediate* category, when the average reproductive activity within a sub-population was centered around the timing of emergence *e*, with a margin of ±0.15 day length.
2. *Dawn-shifted* category, when the average reproductive activity was earlier than the timing of emergence *e*, meaning mating starts happening on the day after the emergence day for those individuals.
3. *Dusk-shifted* category, when the average reproductive activity was later than the timing of emergence, meaning that freshly emerged individuals could start mating on the day of their emergence. Note that within the range of parameter values tested in this study, no such sub-population was observed.

We also characterized the shape of the distribution of reproductive activity timings within each sub-population. Using the *curve_fit* function from the “scipy.optimize” package, we fit a Gaussian curve on these distributions at the end of each simulation. When two sub-populations are detected, we separate our dataset and fit a curve to each of the two sub-populations. We then evaluated the distribution width by using this standardized Gaussian curve, and measured the width at half maximum of the curve (hereafter called average peak width, as seen in fig. 2) to capture the range of *h_a_* values within a subpopulation. This width is strongly correlated to the within-population variance for *h_a_*. Additionally, when two sub-populations were present, we computed the distance between their respective *h_a_* peak.

**Figure 2:**
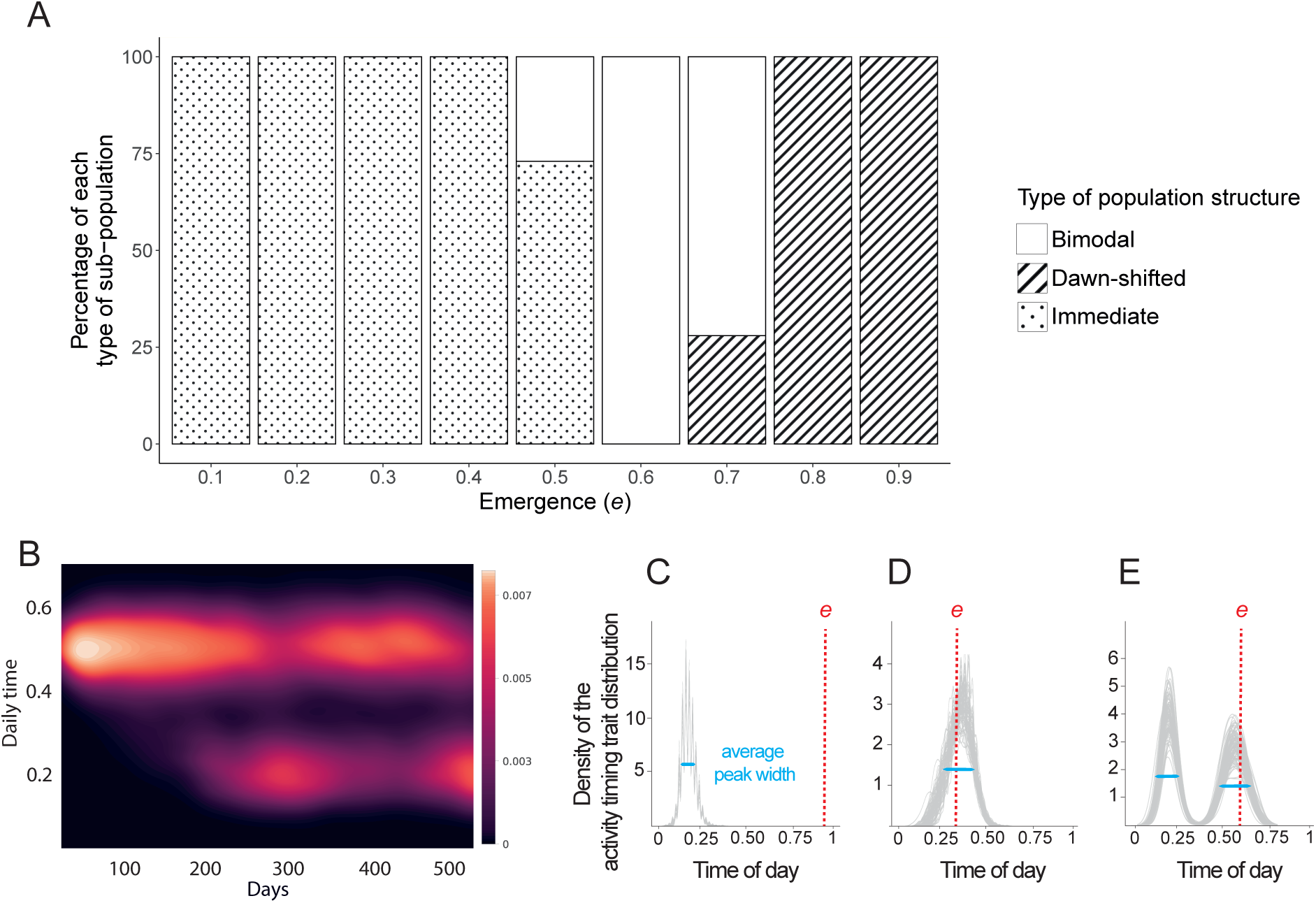
Emergence of sub-populations with different temporal niches depending on the timing of emergence of adults. (A) Percentage of simulations where either one or two subpopulations were observed after 500 days, for 100 replicates per value of the timing of adult emergence *e*. (B) Evolution of the activity timing trait throughout a simulation where bimodality in reproductive timing emerges. (C), (D) and (E) are distributions outcomes of simulations for each type of distribution. Two types of unimodal distribution of the timing of reproductive activity can be observed after 500 days: (C) a single *dawn-shifted* population, where the average time of reproductive activity is below the emergence time *e* or (D) a single *immediate* population. Bimodal distribution of timing of activities leading to the co-existence of two sub-populations can also be observed (E), with both *dawn-shifted* and *immediate* sub-populations. The red dotted lines in (B), (C), (D) and (E) show the emergence timing in our examples (*e* = 0.5, 0.9, 0.3 and 0.6 respectively), and the blue line the average peak width. All simulations were run assuming the same values in the other parameters: *β* = 3, *δ_c_* = 0.1, *G* = 1, *v_e_* = 0.05, *p* = 1, *K* = 1000.

These metrics provide a basis for determining whether two sub-populations are well differentiated or if their reproductive activity timings exhibit a major overlap.

When differentiated sub-populations were observed at the end of a simulation, we quantified the level of genetic differentiation between the two sub-populations, using the *F_ST_* values computed from genetic variations observed at the genetic incompatibility loci. The *F_ST_* is then calculated according to the following equation, from Hudson et al., 1992:

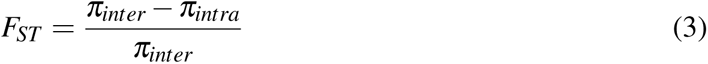

With *π_intra_*and *π_inter_*the average number of pairwise differences between two individuals of the same/different sub-population respectively

Note that under the assumption that intra-population variations are always smaller than inter-population variations, 0 ≤ *F_ST_* ≤ 1, *F_ST_*= 1 is the maximum level of genetic differentiation between the two sub-populations. When differentiated sub-populations were observed at the end of a simulation, we also estimated the level of linkage disequilibrium (*LD*) among loci of the genetic incompatibility loci, among the loci controlling reproductive activity timing *h_a_*, as well as between the two types of loci, using the correlation coefficient between pairs of loci *r*^2^, described as:

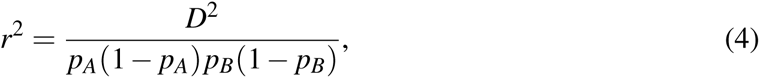

where for an allele *A* of a loci and an allele *B* of a different locus, with respective frequency *p_A_* and *p_B_*, we have *D* = *p_AB_* − *p_A_ p_B_*. Departure of the *r*^2^ value away from zero indicates linkage disequilibrium between the two loci tested. Statistical tests of the *p*-values obtained for the *r*^2^ values computed for the different pairs of loci then rely on Bonferroni correction.

## Results

### Evolution of sub-populations with earlier reproductive activity

Assuming that females are able to mate at maximum once throughout their life (*p* = 1), and assuming no genetic incompatibilities (*G* = 1), our simulations result in either a unimodal or a bimodal distribution of the timing of reproductive activities, depending on the timing of emergence of adults *e* assumed (fig. 2).

- When a unimodal distribution of activity was observed, the average timing of reproductive activity usually follows the timing of emergence for most *e* values (as being active when new virgin individuals emerge is very advantageous, fig. S1A), and a single *immediate* population was thus observed. When the emergence occurs late during the day (*e* ≥ 0.7), individuals are not active close to the emergence time happening at dusk, but rather at dawn (*i.e.* single *dawn-shifted* population). Since our model assumes no mating during the night, late reproductive periods are probably counter-selected. Late timing of adult emergence may thus favor the evolution of reproductive activities early in the morning.
- When a bimodal distribution in the timing of activity is observed, two sub-populations with different temporal niches (*dawn-shifted* niche and *immediate* niche) can co-exist. The subpopulation with the *dawn-shifted* niche has an average timing of reproductive activity close to the start of the day, while the sub-population with an *immediate* niche displays an average timing of activity close to the moment of adult emergence, where more receptive females are available (fig. S1B).

Characterisation of the distributions of the reproductive activity time *h_a_* revealed that when assuming no genetic incompatibilities, the average distance between peaks (when in a bimodal case) in a day was around 0.3 day length (fig. S2A) while the average peak width ranged from 0.16 to 0.19 day length (fig. S2B). Our differentiated subpopulations thus display only a slight temporal niche overlap. When adult emergence occurs very early or very late in the day, the bimodal distribution of reproductive activity disappears (Fig. 2). However, the resulting evolutionary outcomes differ between these two cases. For early emergence (*e <* 0.5), reproductive activity can begin on the same day, favoring the evolution of immediate temporal niches that allow males to mate rapidly with newly emerged virgin females. The absence of available time before emergence constrains the evolution of even earlier activity, preventing further temporal divergence. In contrast, when emergence occurs late in the day (*e >* 0.7), selection favors a single dawn-shifted population, since males with *immediate* timing of reproductive activity would have little mating opportunity before dusk, individuals active early the following day are more likely to encounter virgin females that emerged the previous evening. Overall, these results indicate that temporal niche differentiation is promoted by intermediate emergence times, which provide sufficient temporal space for divergence, whereas extreme emergence times constrain reproductive timing and prevent the evolution of differentiated temporal niches.

We subsequently tested the effect of the variability in the timing of adult emergence (*v_e_*) throughout the day, as it can impact the probability of encounter between males and newly emerged receptive females. Emergence of differentiated sub-population was favored by intermediate values of *v_e_* (fig. S3), while low and high *v_e_* values resulted in unimodal distributions respectively (*immediate* and *dawn-shifted* respectively). While some variability was essential for individuals to diverge in their activity patterns from the emergence peak timing, completely uniform emergence (*i.e.* high *v_e_* values) within a single day prevented the development of distinct temporal niches. See Appendix A for a more in depth analysis of the effect of *v_e_*. The level of variation in the timing of adult emergence within a population thus played a key role in shaping the evolution of temporal niches.

### Behavioral male-male competition and competition for receptive females promoting the evolution of differentiated sub-populations

To test how much the density-dependent male-male competition generates negative frequency dependent selection on the timing of male sexual activity, we compared the proportion of simulations where a bimodal distribution of the timing of activities was observed at *N* = 500 days, when assuming different levels for the cost of behavioral male-male competition. To test for the effect of operational sex-ratio on negative density-dependent selection, we also compare simulations allowing either a maximum of one mating event per female (*p* = 1) *vs.* multiple mating events per female (from *p* = 2 to *p* = 5), using *δ_c_* = 0.1 (fig. 3B). We indeed observe the effect of asymmetry in mating opportunity between males and females in the evolution of differentiated temporal niches: when the maximum number of partners possible for a female increases, the proportion of simulations with co-existing differentiated sub-populations drastically decreases, from one third of simulations when *p* = 1, to being never observed when *p* ≥ 3. The mating asymmetry alone is thus sufficient to generate negative frequency-dependent selection on the timing of reproductive activities through male-male competition. We hypothesize that this competition is driven by a limited availability of females for mating, inherently creating male-male competition for access to females. This competition, in turn, fosters negative frequency-dependent selection, favoring reproductive activity at times when fewer males are competing simultaneously. Note that, in practice, this selection is asymmetric, as only earlier activity timings are selected, as individuals are more likely to encounter virgin females by being active earlier rather than later. Female mating limitations thus strongly promotes the emergence of bimodal distributions in the reproductive activity timing of individuals, without any ecological specialisation or post-zygotic barriers to gene flow.

**Figure 3:**
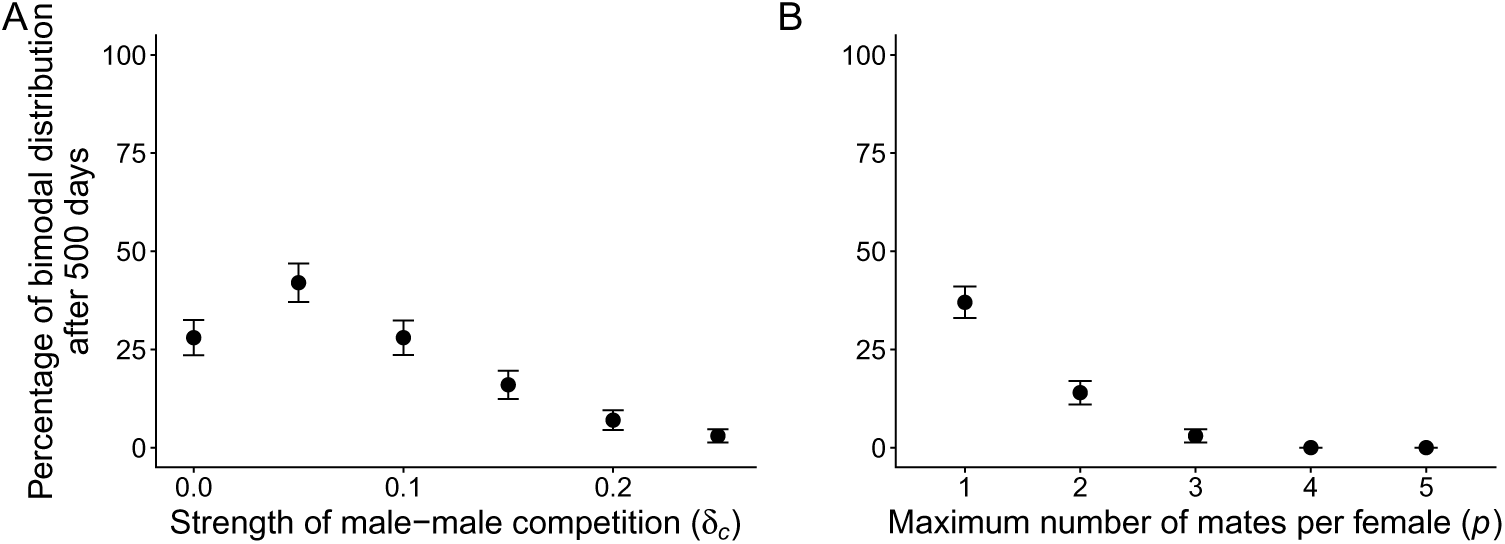
Effects of the male-male competition and male-female asymmetry on the evolution of sub-populations with different temporal niches. (A) Effect of the intensity of male-male competition (*δ_c_*) on the percentage of simulations where a bimodal distribution was observed after 500 days, for 100 replicates per condition. Intermediate values (around *δ_c_* = 0.05) lead to the higher percentage of simulations with coexistence of *dawn-shifted* and *immediate* sub-populations. (B) Effect of maximum mating events per female on the percentage of simulations where a bimodal distribution was observed after 500 days for 100 replicates per condition. Error bars show the standard deviation (SD) computed over the 100 replicates per value of each parameter. All simulations were run assuming the same values in the other parameters *δ_c_*= 0.1, except in 3A, *β* = 3, *G* = 1, *e* = 0.5, *v_e_* = 0.05, *p* = 1, except in 3B, *K* = 1000.

When testing how much behavioral male-male competition affects temporal differentiation, we nonetheless observe an increase in the probability of differentiation into two sub-populations when low competition exists (0.05 ≥ *δ_c_* ≥ 0.1), compared to simulation without competition. However, when the competition becomes too intense (*δ_c_ >* 0.1), the probability of observing differentiated populations decreases, reaching lower values than in simulations without competition. Such decrease may be explained by multiple factors: (1) **Competitive exclusion between the two subpopulations** (Fig. 3A): under weak competition (*δ_c_* = 0.05), the mean duration of coexistence between subpopulations is greater (Fig. S4B) than in simulations without competition (*δ_c_* = 0). However, coexistence duration declines as the intensity of male–male competition increases, indicating that temporal niche differentiation becomes increasingly unstable for certain parameter values. When competition is sufficiently strong (*δ_c_ >* 0.2), coexistence became shorter than in simulations assuming no competition. (2) As competition increased, the variance and average peak width of the *h_a_* distributions in unimodal populations also increased (fig. S5). **Wider distributions of the ancestral population** might prevent the emergence of dawn-shifted sub-populations, as the temporal space is already taken. (3) We additionally tested whether the **mutational effect size of** 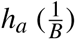 affected the probability of differentiation in high competition settings. By increasing this effect size (*i.e.* a mutation at a loci would result in a higher change in *h_a_* value), the percentage of differentiation at high behavioral male-male competition value sharply increased (from <10% to 40%, see Fig. S6). This suggests that, while selection pressure to be active early is still present even at high *δ_c_* values, the small mutational effect size was preventing populations of early individuals from effectively shifting their activity timing fast enough to not be subject to the extreme competition pressure of the ancestral population, as individuals instead died without mating more frequently. With higher mutational effect sizes, individuals were more likely to shift faster and farther enough in the day to suffer from lower competition pressures, leading to an overall higher differentiation probability.

We thus identify two forms of male-male competition fuelling temporal segregation, in the absence of post-zygotic barriers or ecological specialisation: indirect competition for access to receptive females for mating, as well as direct, density-dependent competition between males (*e.g.* behavioral interactions between males).

### Genetic differentiation favored by competition and genetic incompatibilities

We then explore the effect of genetic incompatibilities on the evolution of differentiated subpopulations with divergent distributions in the timing of reproductive activities, to test for the interaction between pre and post-zygotic factors on population differentiation. We thus investigate the effect of the threshold of genetic divergence at the genetic incompatibility locus *G*, above which reproduction between pairs of individuals becomes unsuccessful. Fig. S7 shows that *G* has a strong effect on the probability of evolution of differentiated sub-populations with divergent temporal niches:

- When the threshold *G* is low (*i.e.* when few allelic variations between parents are sufficient to prevent the generation of viable offspring/imago), the evolution of differentiated sub-populations is relatively limited. Such drastic incompatibility threshold may strongly limit offspring viability and thus frequently prevent the evolution of unusual timings of reproductive activity.
- When assuming intermediate values (0.4 ≤ *G* ≤ 0.5), the percentage of simulations where we observe the evolution of differentiated sub-populations becomes much higher (*>* 98%), but decreases when the threshold further increases.

The evolution of differentiated populations is thus more likely to happen for moderate values of the threshold *G*, where post-zygotic barriers can emerge between sub-populations, but do not prevent differentiation, as viable offsprings can still be produced from crosses between individuals belonging to the two emerging sub-populations. The level of genetic differentiation between these sub-populations can then be estimated using the fixation index (*F_ST_* ).

When assuming genetic incompatibilities (0 *< G <* 1), the *F_ST_*values are higher when *G* is smaller (fig. 4A). Low values of *G* strongly promote assortative mating between individuals with similar genotypes, likely driving the genetic differentiation between *dawn-shifted* and *immediate* sub-populations. When the threshold *G* becomes even higher (*G* ≥ 0.6), *F_ST_* values become low, coinciding with the decrease in the probability of population differentiation shown in fig. 4.

**Figure 4:**
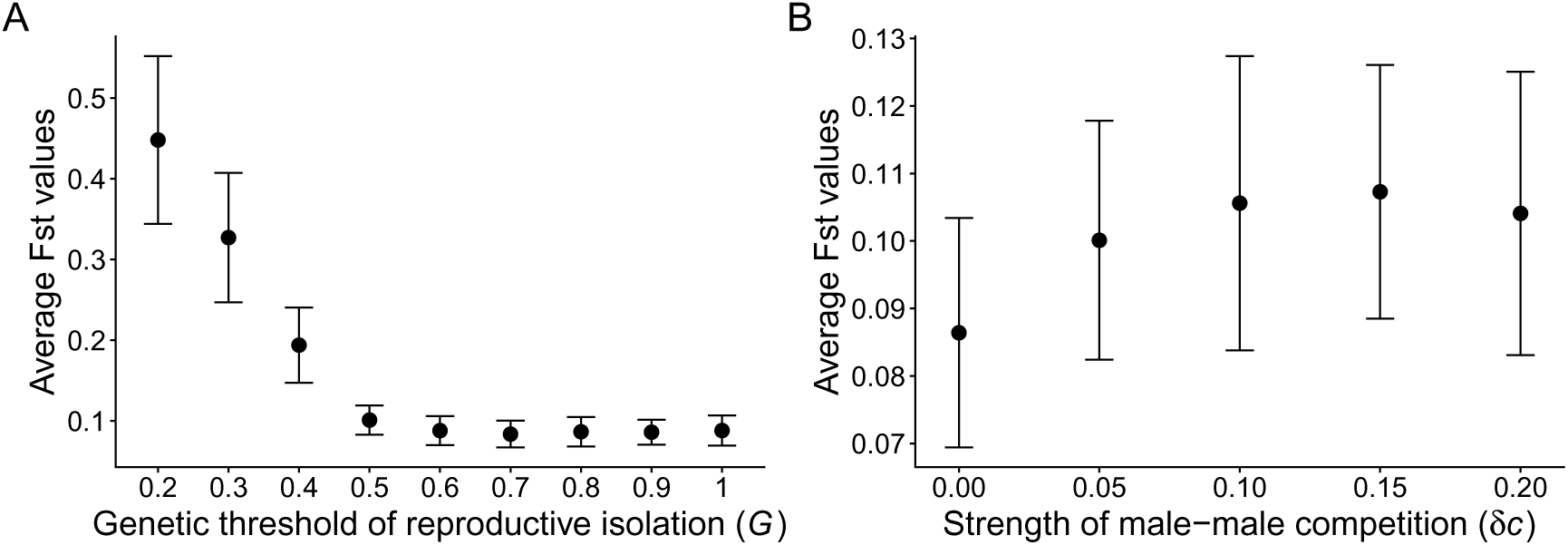
Effect of genetic incompatibility and male-male competition on the genetic differentiation between sub-populations with different temporal niches. The average *F_ST_*between sub-populations where a bimodal distribution was observed after 500 days, assuming genetic incompatibilities is shown, for 100 replicates per values. (A) Effect of the proportion of loci *G* required for mating incompatibilities on the level differentiation between sub-populations, estimated by *F_ST_* . Small and moderate values of *G* promote high *F_ST_* values between sub-populations. (B) Effect of male-male competition intensity *δ_c_*on the *F_ST_* values between differentiated populations. When the male-male competition intensifies, average *F_ST_* between sub-populations increases. Error bars show the SD computed over the 100 replicates per value of each parameter. All simulations were run assuming the same values in the other parameters: *β* = 3, *δ_c_* = 0.1, *G* = 0.5 for fig. 4B, e = 0.5, *v_e_* = 0.05, *p* = 1, *K* = 1000.

When studying the effect of the genetic incompatibilities promoting the co-existence of the two sub-populations, we found that the duration of the co-existence of the two sub-populations was maximal when *G* = 0.5 (fig. S4A). Notably, when *G >* 0.5, coexistence was much shorter, which probably explain why fewer differentiated sub-populations are observed at the end of simulations in these cases, as they struggle to maintain themselves. In extreme cases (*G* = 0.1), every simulation lead to the extinction of the whole population, because most individuals failed to generate any offspring (fig. S7), while for values of *G >* 0.2, no extinction is observed. While the effect is weak, stricter threshold (*i.e.* lower *G* values), promotes a larger temporal gap in reproductive activity timing between sub-populations (fig. S2A), as well as narrower width of peaks (fig. S2B). Genetic incompatibility thus seem to reinforce the temporal divergence between the two sub-populations, as well as the specialisation into narrower temporal niches.

Furthermore, the strength of male-male competition also had an effect on genetic differentiation between populations (captured by *F_ST_*), with genetic differentiation increasing with the intensity of competition (fig. 4B): *F_ST_* values where higher in simulations with high competition (*δ_c_* = 0.2) as compared to simulations without (*δ_c_* = 0). Thus while high values of *δ_c_* decreases the probability of population differentiation (see fig. 3A), in the fewer cases where sub-populations do evolve (see section above), high levels of genetic differentiation are favored by the intense level of competition. This suggests that genetic drift might also play a role in the evolution of genetic differentiation in this case, with the lower number of males resulting in a decreased gene pool. Overall, by favoring temporal differentiation, the male-male competition can promote genetic divergence between the emerging sub-populations even in the absence of genetic incompatibilities, highlighting the strong effect of pre-zygotic barriers in the differentiation process.

Finally, to investigate the coupling between pre and post-zygotic barriers to gene flow between the two sub-populations, we quantified linkage disequilibrium among genetic incompatibility loci, activity timing loci, and between both types of loci. When computing *r*^2^ in the whole population, we can observe that for lower values of G, *r*^2^ display higher median values, but also higher maximum values (fig. 5). Interestingly, activity timing loci display a relatively constant medium value of *r*^2^ compared to genetic incompatibility loci, which decreases with *G*, showing how selection on activity timing is present even without genetic incompatibilities. Stricter genetic incompatibilities are nonetheless promoting linkage disequilibrium more frequently, as well as more strongly. This is expected, as a stricter threshold for genetic incompatibilities is likely to fix more alleles at the different loci within the two populations. Stricter genetic incompatibilities are thus promoting more frequent linkage disequilibrium between loci acting as post-zygotic barrier and the locus controlling the timing of reproductive activity, acting as pre-zygotic barrier. We also observe *LD* between incompatibility and timing loci for *G <* 0.5. This indicate some level of coupling between pre and post-zygotic mechanisms of reproductive isolation, even though free recombination in our system likely .

**Figure 5:**
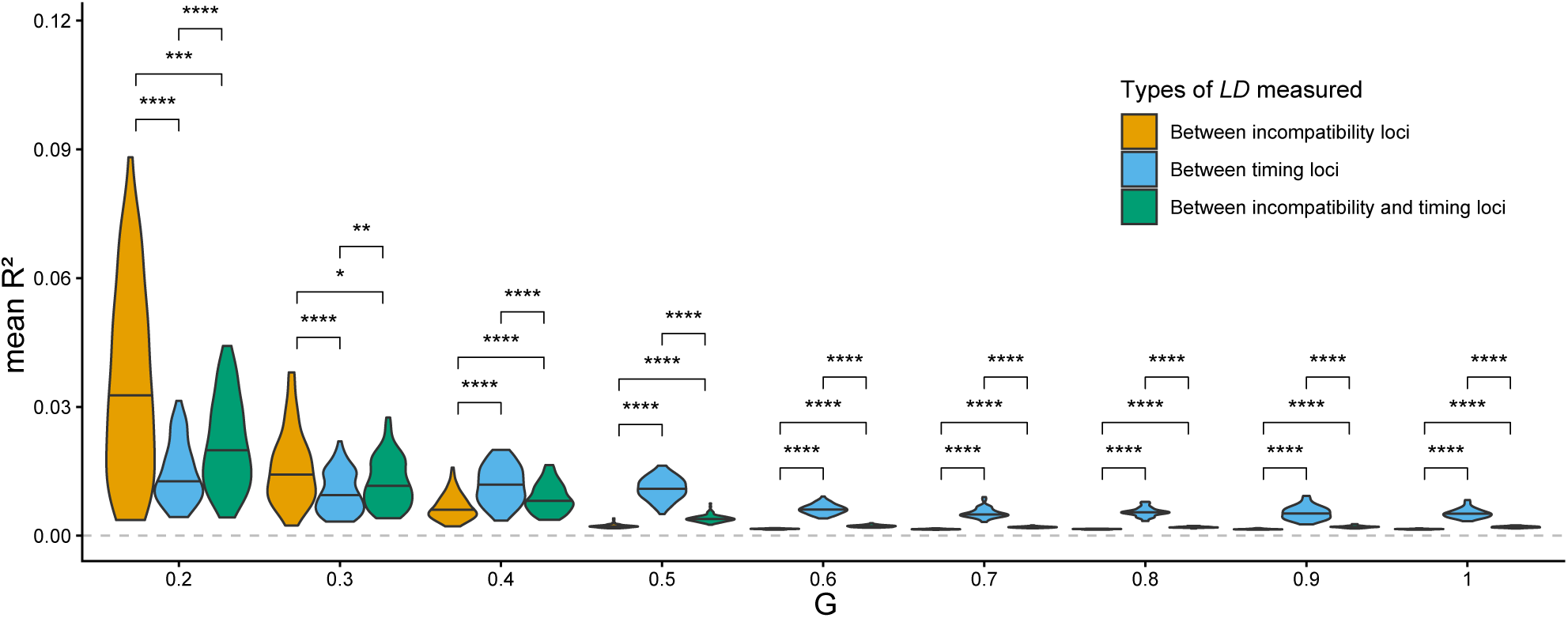
Characterisation of the linkage disequilibrium between activity time and the genetic incompatibility loci, by computing the mean *r*^2^ value. We quantified linkage disequilibrium between genetic incompatibility loci (yellow), activity timing loci (blue), and between both (green). Significant differences between mean *r*^2^ values are indicated. We can observe that for lower values of G, *r*^2^ display higher median values, but also higher maximum values. Stricter genetic incompatibilities are thus promoting stronger linkage disequilibrium between incompatibility loci, as well as between incompatibility and timing loci. All simulations were run assuming the same values in the other parameters: *δ_c_* = 0.1, *β* = 3, *e* = 0.5, *v_e_* = 0.05, *p* = 1, *K* = 1000.

Genetic differentiation between sub-populations thus appears to be positively correlated to both the strength of behavioral male-male competition and the incompatibility threshold between divergent genomes. The linkage disequilibrium between activity timing loci and genetic incompatibility loci generates a coupling of pre and post-zygotic barriers to gene flow between sub-populations that may further promote sympatric speciation.

## Discussion

### Evolution of temporal niches of reproductive activities

Using our model describing the evolution of *daily* reproductive activity niche, we show that subpopulations with different timings of reproductive activities can emerge through time. Because we assumed that offspring have a timing of reproductive activity intermediate between their parents (*modulo* the mutational effect *µ*), our model limits the emergence of temporal differentiation within our simulated populations. Moreover, our allelic structure create a mutational bias in favor of an average phenotype (*h_a_* = 0.5), with establishment of phenotypic values near the boundary [0,1] being unlikely in the absence of evolutionary forces. Despite this reduced opportunity of divergence stemming from the genetic architecture assumed for the control of the timing activity in our model, we frequently observed differentiation into two subpopulations. The differentiation into different temporal niches is mainly due to the limited availability in receptive females, but is also promoted by the costs generated by antagonistic male-male interactions. Negative density-dependent selection acting on the timing of reproductive activities can therefore promote differentiation into different temporal niches. Another model recently investigated the ecological factors promoting daily allochronic speciation in the *Spodoptera frugiperda* moth (Van Doorn et al., 2024), by assuming divergent ecological selection favoring divergent adaptive timings of reproduction in the rice *vs*. corn eating populations. In line with our own results, the operational sex-ratio (controlled in their model by monogamy *vs*. polygamy during the night) also tuned the intensity of male-male competition and the resulting negative frequency-dependent selection acting on the activity timing was a key parameter shaping the evolution of allochrony.

In contrast, our model generates allochrony strictly based on male-male competition for virgin females, without assuming any other ecological mechanisms generating disruptive selection. However, when assuming independent control of the activity timing between males and females, we found that temporal segregation rarely emerged (fig. S8). This suggests that male behaviour indeed strongly drive temporal differentiation in our model. When assuming a common control of reproductive activities in males and females, the emergence of differentiated sub-populations is likely promoted by assortative mating between individuals with similar neutral genomes (Kondrashov and Shpak, 1998), stemming from a synchronicity of reproductive activities. The timing of reproductive activity acts as a grouping magic trait (Kopp et al., 2018), which might be reduced when the timing of activity evolves independently in males and females, therefore restricting the conditions enabling the evolution of differentiated timing of activity (fig. S8). Moreover, we observe linkage disequilibrium between our ecological trait (activity timing) and neutral traits when assuming common control of reproductive activity between males and females. This observation aligns with Dieckmann and Doebeli (1999) prediction that *LD* between neutral and ecological traits facilitates speciation under non-random mating.

In previous models describing the evolution of flowering times in plants, negative density dependent selection stemming from pollinator availability has already been described as involved in population subdivisions into different temporal niches (Devaux and Lande, 2008). Nevertheless, this division was generally found symmetrical around the ancestral flowering time. In our model however, the evolution of differentiated temporal niche is not symmetrical. We indeed observed an asymmetrical distribution in our *daily* model, with new sub-populations having earlier activity timings, due to competition for receptive females. We never observed *dusk-shifted* sub-populations, where individuals would be selected to be active later than emergence in a day. This shows an asymmetric positive selection for earlier *h_a_* values and not for later ones, because of male-male competition to access virgin females as soon as possible during the day. This difference is likely due to the gradual maturation of the different flowers within each plant assumed in Devaux and Lande, 2008: in this model, each individual possess multiple independent flowers becoming open on different days throughout the flowering period. The ovules of the flowers carried by a single plant could then be fertilized at different moments within the activity period. Hence, there is no specific advantage for early pollen as compared to late ones, allowing the evolution of both early and late-shifted flowering periods. This result highlights a potential major difference in terms of allochronic speciation outcomes depending on mating systems.

In perennial species where individuals are potentially available for reproduction all year round, changes towards earlier breeding times during the year is shown to occur in wild populations of *Parus major* (Charmantier et al., 2008). Furthermore, at the daily scale, fitness advantages associated with earlier sexual activity (mate guarding and extra-pair mating) in males were also reported in behavioral experiments carried out in *Parus major* (Greives et al., 2015). These empirical results suggest that changes in the timing of reproductive activities can occur in the wild, because of both plasticity and/or genetic variations, and might be favored by selective pressures. In our model, we assumed that reproductive activity times were independent from the emergence time, which was fixed during our simulations. Nevertheless, the reproductive timing of activity (modelled here as *h_a_*) and the emergence (*e*) are likely co-evolving in natural populations and are probably jointly influenced by the circadian clock, as shown in Drosophila (Calatayud et al., 2007; Kumara et al., 2015). Supplementary simulations allowing the emergence timing to evolve independently of reproductive timing of activity demonstrated the emergence of differentiated sub-populations (fig. S9), both evolving towards earlier activity timings. These results demonstrate the strong effects of negative frequency dependence stemming from male-male competition on sub-population differentiation and coexistence. In the model where *h_a_* and *e* co-evolve, selection on being active as early as possible is strong in males given the absence of a fixed emergence timing during midday. The two sub-populations with earlier activity timings did not merge, even in the absence of genetic incompatibilities. This shows how negative frequency dependence can maintain two populations in sympatry, with separated temporal niches. The evolution of the temporal window of reproductive activity predicted by our model could thus happen in natural populations, in presence of genetic variations controlling for the timing of reproductive behavior, like courtship or mate searching activities.

### Ecological factors promoting temporal divergence between sub-populations

In our model, the coexistence of sub-population was sometimes transient. Indeed, the co-existence of sub-populations varied depending on the strength of male-male competition *δ_c_* and the threshold of genetic divergence *G*: we notably showed that moderate values of *δ_c_*and *G* led to a longer co-existence duration. The limited duration of coexistence may be linked to the absence of joint evolution of other isolating mechanisms, such as host plant specialisation or coupling with abiotic factors. In our model, we indeed investigated the effect of neutral mutations triggering incompatibilities between divergent individuals. Nevertheless, if these mutations accumulating in the sub-populations with different temporal niches were under divergent selection, this may further reinforce co-existence of differentiated populations.

Temporal differentiation can be facilitated by variation in resource availability throughout the day, that may condition the timing of feeding *vs.* reproductive activities. In insects, host shifts may thus enhance the evolution of temporal niches. For instance, variation in the timing of flowering (afternoon *vs*. dusk) was reported among closely-related species of sphingophilous plants of the genus *Hemerocallis* (Hasegawa et al., 2006). Specialisation into these different plant species may induce temporal divergence in pollinators. Multiple other plants also display such daily variations flowering/pollen release patterns (Matsumoto et al., 2015; Stone et al., 1996), that may condition the evolution of daily temporal niche in pollinating insects.

Abiotic factors such as temperature may also enhance allochronic speciation, if the reproductive success is lower when individuals are not in their optimal temperature range (Dolgin et al., 2006; Keller and Seehausen, 2012). Since thermal conditions usually vary throughout the day, the evolution of the temporal niches can be influence by variation in adaptation to thermal conditions. Differences in reproductive success was detected between sister species locally adapted to different thermal environments (Matute et al., 2009), probably as a by-product of divergent adaptation. As variations in selection of thermal resistance hinder local adaptation, species are likely to be adapted to a relatively small range of temperature conditions (Kawecki and Ebert, 2004), and variations in thermal niches can be a strong barrier to reproduction. These temperature fluctuations are shown to be a source of diversification among species (Kleckova et al., 2015), with empirical data on animal and plant species (Keller and Seehausen, 2012). The evolution of divergent thermal tolerance could co-evolve with the evolution of divergent temporal niches, at both the seasonal and daily scales.

### Allochrony as a reinforcement mechanism in the divergence of species

Allochrony could also act as a reinforcement mechanism, when hybrids are already counter-selected because of post-zygotic barriers. In Butlin and Smadja, 2018, such coupling between pre and post-mating barriers was shown to evolve, but disentangling whether allochrony is the driving force of speciation or just stem from reinforcement remains challenging (see Taylor and Friesen, 2017).

To study the emergence of reproductive barriers in our population, we observed population dynamics with and without post-zygotic barriers to reproduction, represented by non-viable zygotes when the parents were genetically differentiated. When those genetic incompatibilities were introduced, the evolution of sub-populations with different temporal niches was more likely to happen for certain values of the genetic threshold *G*. More importantly, genetic incompatibilities fuel genetic differentiation between sub-populations, as the gene flow between sub-population keeps occurring in absence of these genetic incompatibilities. Potential sympatric speciation could thus be driven by incompatibility selection (Artzy-Randrup and Kondrashov, 2006), even in the absence of other forces promoting ecological divergence. By testing the proportion of loci *G* required for incompatibilities, we found that intermediate values seemed to promote the longevity of small-sized *dawn-shifted* sub-populations. These values seem to allow a low level of gene flow that promoted co-existence, while allowing genetic differentiation between sub-populations. This is consistent with simulations for neutral speciation by distance (Baptestini et al., 2013, de Aguiar et al., 2009), showing similar results when incorporating a genetic mating distance *G*, with intermediate values of maximum genetic distance for reproduction producing higher counts of species. Temporal divergence could thus act as a coupled mechanism to other differentiation factors, by limiting gene flow between sub-populations.

Furthermore, during the differentiation into different temporal niches, offspring with intermediate temporal niches could suffer from elevated competition costs, as they would pay costs from negative interactions with individuals from both parental temporal niches. These intermediate offspring also have a low chance of occurring when genetic incompatibilities are added, since those two sub-populations are usually genetically differentiated, and hybrids were considered inviable. According to simulations performed by Liou and Price, 1994, having inviable hybrids can lead to higher rates of speciation than with infertile ones, because the lack of individuals with intermediate phenotypes reduces competition (for resources/mating partners) for both sub-populations. This was also shown to make extinction less likely for sub-populations, thus promoting coexistence between divergent populations. This coexistence is thus likely conditioned by the availability of empty temporal niches. Indeed, our simulations showed that a certain width of temporal space is necessary for sub-populations to emerge earlier within the day, as they otherwise are too close to the alternative population and male-male competition may then lead to competitive exclusion. Smaller niches (*e.g.* reduced variance of activity timing within emerging sub-populations) would result in more temporal space available for different sub-populations to emerge, and thus provide more opportunities for speciation (Sexton et al., 2017). We indeed observed that temporal niches were narrower (the average peak width of sub-populations decreased) and further apart when implementing stricter threshold for genetic incompatibilities (fig. S2), a behavior akin to reinforcement. Niche breadth is generally shown to be an important factor influencing speciation (Hardy and Otto, 2014) and this effect is confirmed in our study, showing that gradual evolution of divergent temporal niches can be promoted by post-zygotic incompatibilities. While this is true at the population level, we also ran additional simulations with varying values of the individual patrolling timing *w_a_*. We found that with shorter times of reproductive activity, bimodality was not promoted, and we rather observed more uniform distributions (greater width of activity peaks, Fig. S10B). We believe that this is due to an effective decrease in male-male competition, since shorter activity periods reduce the number of males and females a male can encounter. Similarly, longer periods of activity did not promote bimodality, but rather *immediate* populations (Fig. S10B): early sub-populations then strongly overlap with the ancestral one, and thus being active early or late presented no particular advantage. Thus, pre-zygotic factors such as individual length of reproductive timeslot might also play a role in allochronic speciation.

Our study therefore highlights the temporal variation in reproductive activity as an important component of the ecological niches, potentially reinforcing mating barriers between diverging subpopulations. We moreover show that male-male competition alone can initiate population divergence in a sympatric setting, without any ecological trait divergence or post-zygotic isolation.

## Data availability

The data underlying this article are available in the Dryad Digital Repository, at: http://datadryad.org/share/VMLo_jU1B507QvxuZorosNRw_NVU1uuLoSW62d5YBIk

The data underlying this article are available in the Zenodo Digital Repository, at : https://doi.org/10.5281/zenodo.18267436

## Author contributions

Titouan BOUINIER conceived the experiments, collected the data, and wrote the original draft. Arthur BRUNAUD developed the first versions of the code. Titouan BOUINIER, Violaine LLAU-RENS and Charline SMADI reviewed and edited the writing at all stages of revision.

## Funding

This study was funded by the European Union (ERC-2022-COG - OUTOFTHEBLUE - 101088089). Views and opinions expressed are however those of the authors only and do not necessarily reflect those of the European Union or the European Research Council. Neither the European Union nor the granting authority can be held responsible for them.

## Conflict of interest

None.

## Acknowledgments

We thanks Amaury LAMBERT and Guillaume ACHAZ for their thoughtful contributions to the code and interesting questions.

## Notes

http://datadryad.org/share/20xSLjxYGxlkU1QV-vTMITztq4XeQ427NGphEm-mNKg

### Competing Interest Statement

The authors have declared no competing interest.

### Summary of Updates

Updated Methods and Results, Figure 1,2 and 5 revised.

