## SupplementaryMaterial for "Evolution of divergent daily temporal niches shaped by male-male competition can generate sympatric speciation" (from another version of this preprint: version 4, 2025-08-04)

Supplementary Material :

**APPENDIX A: Details on model**

The day (a continuous interval from sunrise (*t* = 0) to sunset (*t* = 1)) is split into smaller time intervals at the start of the day. These intervals are defined by the temporal position of *known* events.

- *Known* events are the transitions from resting to reproductive activity (and vice versa) for every individual, as well as the emergence of new adult individuals. The number of known events is thus at least 2*Number of individuals + Number of emerging individuals.

A second type of stochastic event can happen between two *known* events, which are *unknown* events.

- *Unknown* events are dependent on the state of the population at a certain moment in time, such as mating or death events. The individual involved in an event is randomly chosen, uniformly among individuals exhibiting the same status. The time at which those events happen is drawn in an exponential distribution, based on the current numbers of active males **N_a_^♂^**, resting males **N_r_^♂^** , available active females **N_a_**^♀^, resting available females **N_r_**^♀^, and unavailable females **N_m_**^♀^. We continue drawing new mating or death events until the drawn time exceeds the time of the next known event. At that point, the known event is prioritized and takes place instead.

*Unknown* events occurrence times and rates are computed using the instantaneous mating rate λ_r_ and the instantaneous death rate λ_d_, given by:

(1)

λ_r_ = β * **N_a_^♀^** ^*^ **N_a_^♂^**

(2)

λ_d_ = d * (**N_a_^♂^** + **N_r_^♂^** + **N_a_^♀^** + **N_r_^♀^** + **N_m_**^♀^) + δ_e_ * (**N_a_^♂^** + **N_a_^♀^**) + δ_c_ * **N_a_^♂^** * (**N_a_^♂^**-1) + δ_l_ *$\sum_{i=0}^{N_{m}^{\mathbf{♀}}} E_{i}$

The time at which those events happen is drawn in an exponential distribution of parameter $\frac{1}{\lambda r+\lambda d}$. If this occurring time is smaller than the occurring time of the next known event, then a mating or a death event happens. If this occurring time is higher than the occurring time of the next event, nothing happens except the next known event. To determine which event between mating and death occurs, a value is drawn from a uniform distribution *U*(0,1) and is compared to $\frac{\lambda r}{\lambda r+\lambda d}$. If the value drawn is smaller, then a mating event occurs. If not, we compare it with $\frac{\lambda r}{\lambda r+\lambda d}$. + $\frac{d* N_{r}^{\mathbf{♂}}}{\lambda r+\lambda d}$, which corresponds to the death of an inactive male. If the value is still greater, we compare with $\frac{\lambda r}{\lambda r+\lambda d}$. + $\frac{d* N_{r}^{\mathbf{♂}}}{\lambda r+\lambda d}$ + $\frac{(d + \delta_{e}\mathbf{)}* N_{a}^{\mathbf{♀}}}{\lambda r+\lambda d}$ (death of an available active female), and so on.

In the case of unavailable females, we consider the number of eggs $E_{i}$ laid by each individual.

As such, there are **N_m_**^♀^ events we need to check:

We compare the value drawn from the uniform distribution to with:

- $\frac{\lambda r}{\lambda r+\lambda d}$ + … +$\frac{d+ \delta_{l} * E_{1}}{\lambda r+\lambda d}$ for the death of unavailable female number 1;
- $\frac{\lambda r}{\lambda r+\lambda d}$ + … + $\frac{2d+ \delta_{l} *(E_{1}+E_{2})}{\lambda r+\lambda d}$ for the death of unavailable females number 2;

and so on until the event of the death of unavailable female number **N_m_**^♀^:

- $\frac{\lambda r}{\lambda r+\lambda d}$ + … +$\frac{(N_{m}^{\mathbf{♀}}* d)+ \delta l *\sum_{i=1}^{N_{m}^{\mathbf{♀}}} E_{i}}{\lambda r+\lambda d}$.

**Variation in the timing of adult emergence shapes population differentiation**

Because the variability in the timing of adult emergence *v_e_* throughout the day can impact the probability of encounter between males and newly emerged receptive females, we run simulations for different levels of variability in the timing of emergence, assuming that females can only mate once (*p* = 1) and no genetic incompatibilities, and followed the distribution of the activity timing trait *h_a_* through time.

Assuming that adults emerge around noon (*e* = 0.5), we run 100 simulations for each level of variation in the timing of the emergence and observed three main evolutionary outcomes (fig. S9):

- When assuming low variability in the emergence timing (*v_e_* < 0.10), a unimodal distribution of timing of reproductive activity is observed in a majority of simulations, with a single *immediate* population. The low variance in the timing of sexual activity in receptive females means that males are strongly selected to be active at this moment, therefore limiting the opportunities for temporal differentiation.
- When assuming moderate levels of variations in the emergence timing (0.05 > *v_e_* > 0.09), up to a quarter of simulations result in a bimodal distribution of the sexual activity timing, generating sub-populations with different timings of reproduction, with a *dawn-shifted* sub-population co-existing with an *immediate* sub-population.
- When assuming high variability in the emergence timing (*v_e_* > 0.24), a unimodal distribution of timing of reproductive activity is observed, with sexual activities occurring early in the morning (*dawn-shifted* population), away from the mean emergence time (*e* = 0.5). The high uncertainty in the timing of availability in receptive females likely promotes the evolution of the earliest possible timing of male reproductive activities. The even temporal distribution of receptive females throughout the day no longer promotes sexual activities around the emergence peak, and only the advantage associated to be the first one to catch an receptive female remains in males. Male-male competition likely generates a race promoting earlier male reproductive activities, leading to the persistence of a single *dawn-shifted* population and limiting temporal differentiation.

Variation in the timing of adult emergence within a population thus plays a key role in shaping the evolution of temporal niches. While some variability is essential for individuals to diverge in their activity patterns from the emergence peak timing, completely uniform emergence within a single day prevents the development of distinct temporal niches.

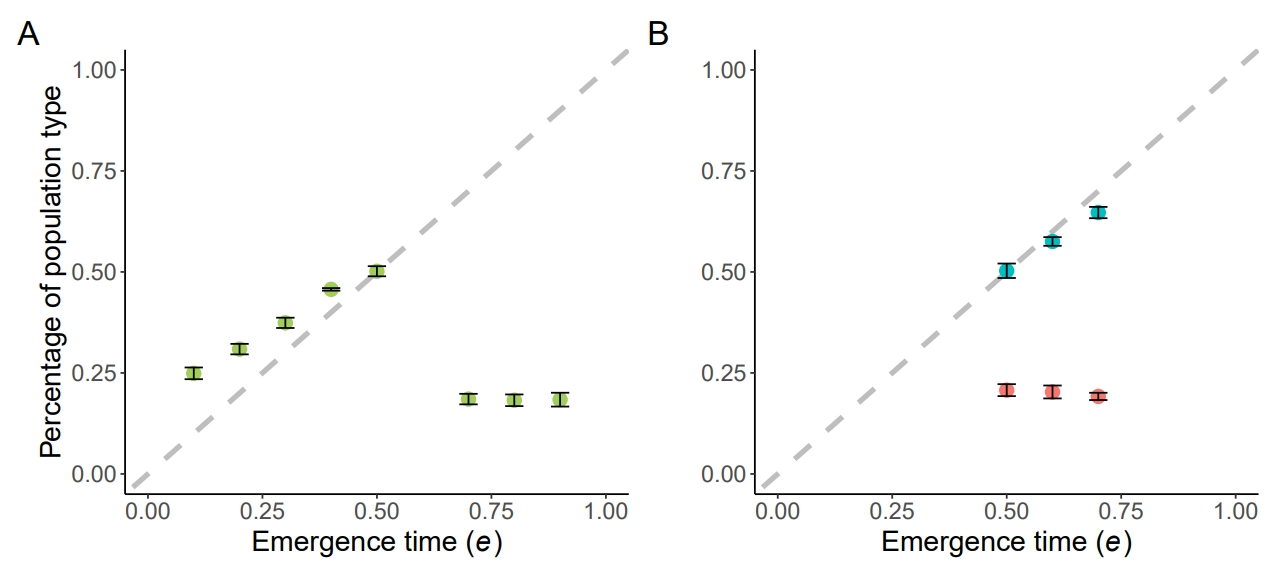

Figure S1: **Temporal niches in the sub-populations, depending on the timing of adult emergence**, estimated by the average reproductive activity timings, obtained in simulations assuming different values of the timing of emergence of adult e. We run 100 simulations per condition, and distinguished them based on the distribution of the timing of reproductive activity *h_a_* after 500 days. We separated outputs with distributions that were either unimodal (A) or bimodal (B). In the bimodal case, dawn-shifted and immediate temporal niches can be observed, where the average *h_a_* within the sub-populations are shown in red and blue respectively. Dashed lines then show the emergence time e in the corresponding simulations.

Error bars represented by one SD. All simulations were run assuming the same parameter values: δc = 0.1, β = 3, G = 1, Ve = 0.05, p = 1, K=1000.

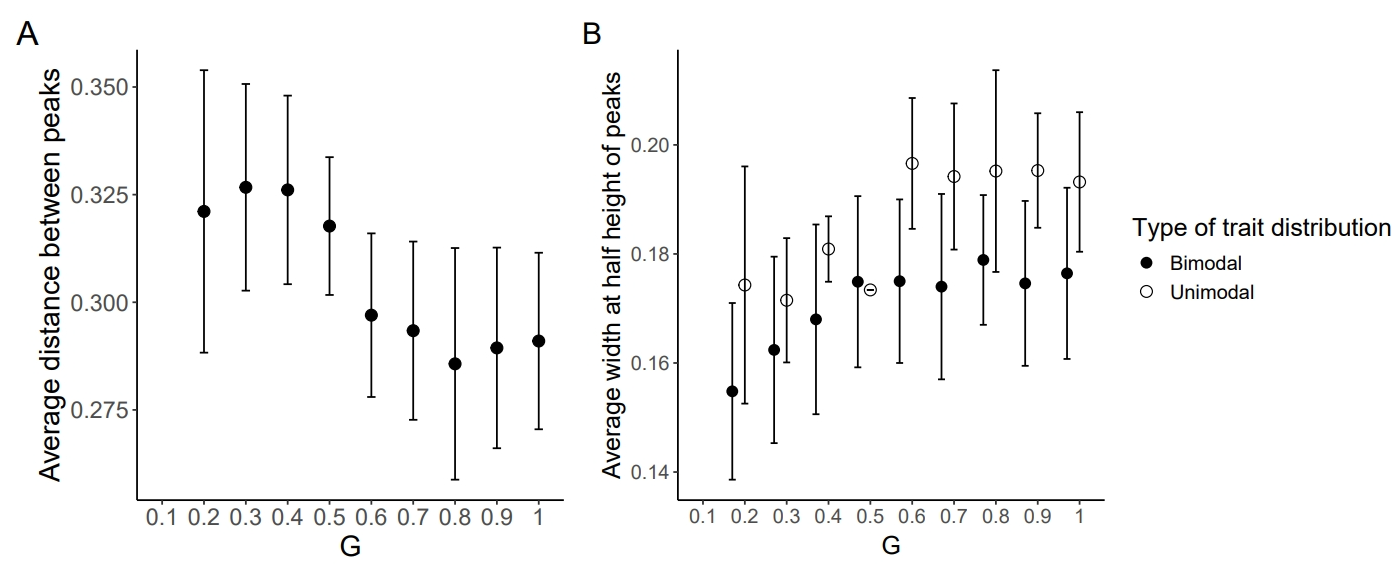

Figure S2: **Peaks characterization via distance between peaks and width of peaks, depending on the proportion of loci necessary to generate incompatibilities.** We ran 100 simulations for each parameter, assessing the (A) average distance between peaks at the end of simulations, when two peaks were detected. We also measured (B) the average width of the peaks at half height, using a gaussian fit for each peak. This characterization ensured that our peaks did not overlap, and were far enough apart to be considered two differentiated sub-populations. Notably, in bimodal cases, activity timing peaks were slightly narrower than in unimodal cases.

Error bars are one SD. All simulations were run assuming the same parameter values: δc = 0.1, β = 3, e = 0.5, Ve = 0.05, p = 1, K=1000.

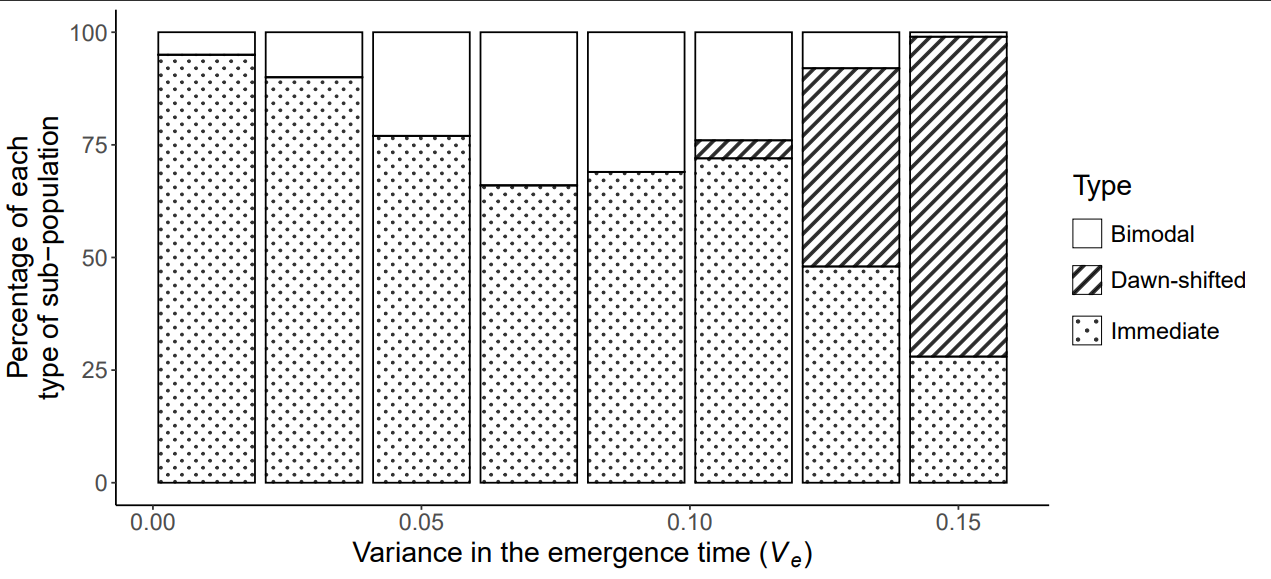

Figure S3: **Emergence of sub-populations with different temporal niches depending on the variance of the adult emergence timing.** Percentage of simulations where either one or two sub-populations were observed after 500 days, for 100 replicates per value of the timing of adult emergence.

All simulations were run assuming the same parameter values: δc = 0.1, β = 3, G = 1, e = 0.5, p = 1, K=1000.

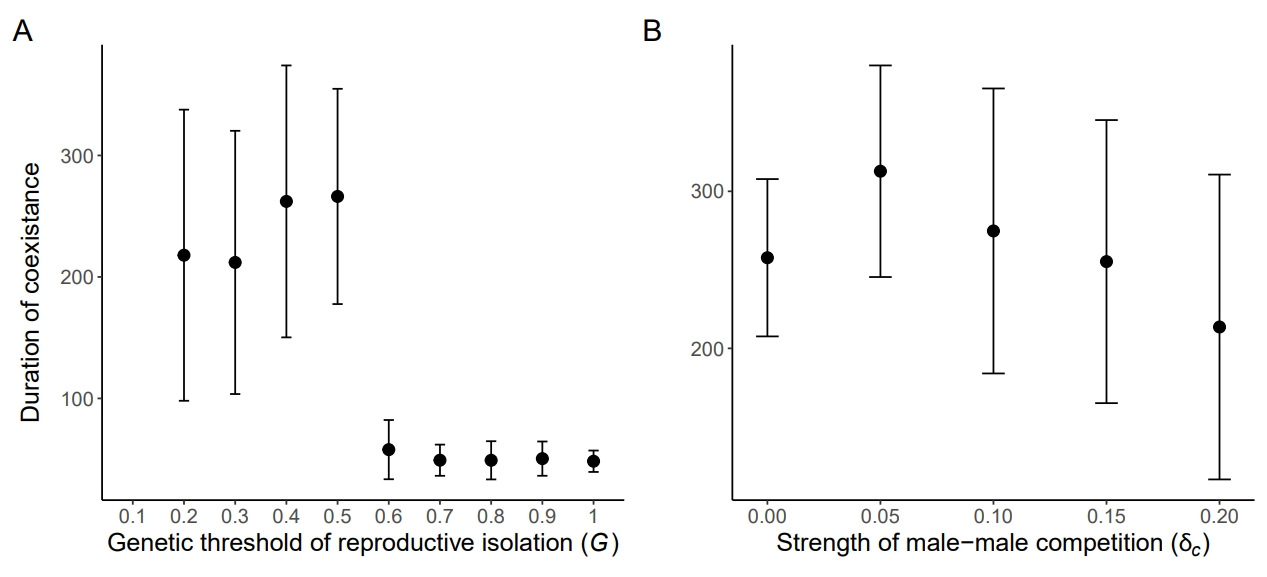

Figure S4: **Duration of coexistence between dawn-shifted and immediate sub-populations.** We ran 100 simulations per parameter, for (A) the proportion of loci necessary to generate incompatibilities, and (B) behavioral male-male competition.

Error bars are one SD. All simulations were run assuming the same parameter values: δc = 0.1, β = 3, e = 0.5, Ve = 0.05, p = 1, K=1000.

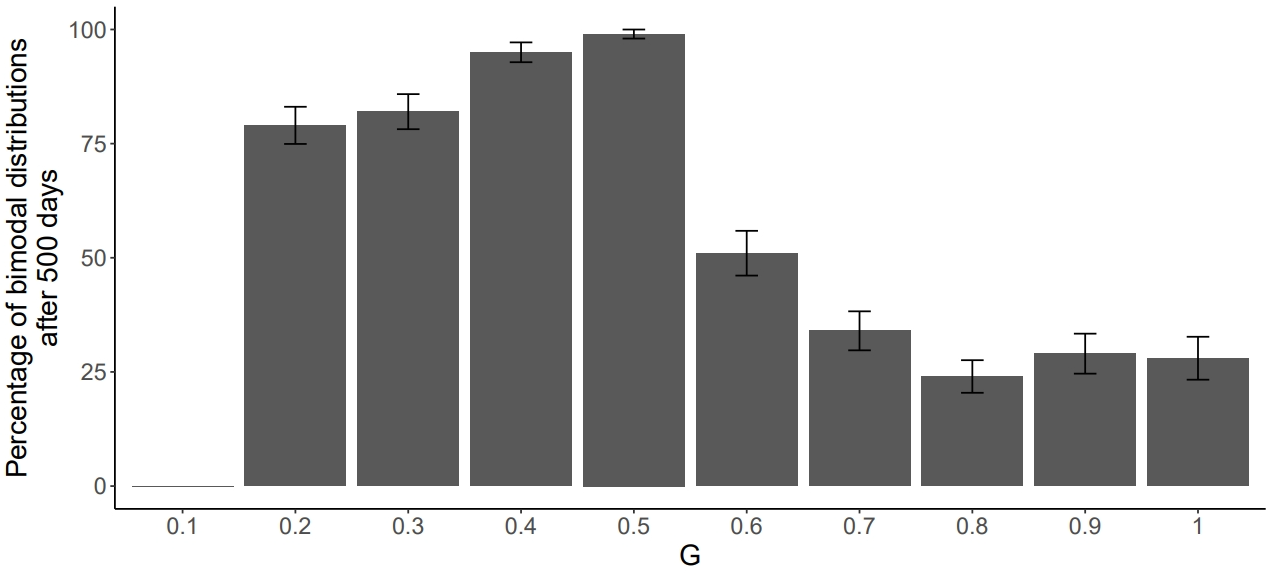

Figure S5: **Effect of the proportion of loci required to generate incompatibility (*G*) between individuals on the evolution of sub-populations with different temporal niches.** Percentage of simulations where a bimodal distribution in the timing of activities is observed at 500 days, depending on the proportion of loci G necessary to trigger incompatibility.

Error bars are one SD. All simulations were run assuming the same parameter values: δc = 0.1, β = 3, e = 0.5, Ve = 0.05, p = 1, K=1000.

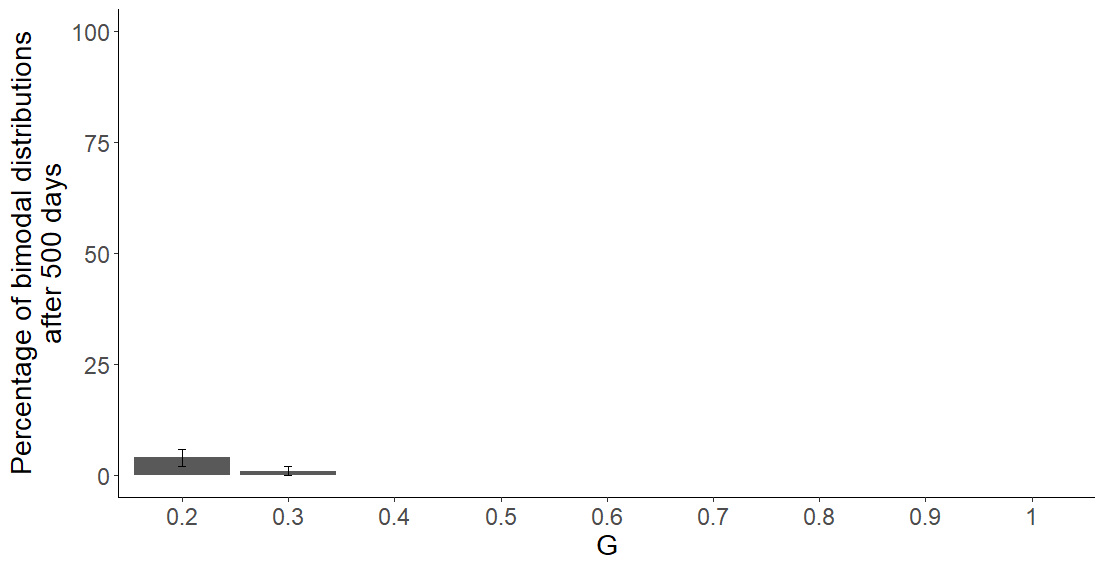

Figure S6 : **Effect of the independent evolution of sexual activity timings in males and females.** Percentage of simulations where a bimodal distribution in the timing of activities is observed at 500 days, depending on the proportion of loci G necessary to trigger incompatibility.

Error bars are one SD. All simulations were run assuming the same parameter values: δc = 0.1, β = 3, e = 0.5, Ve = 0.05, p = 1, K=1000.

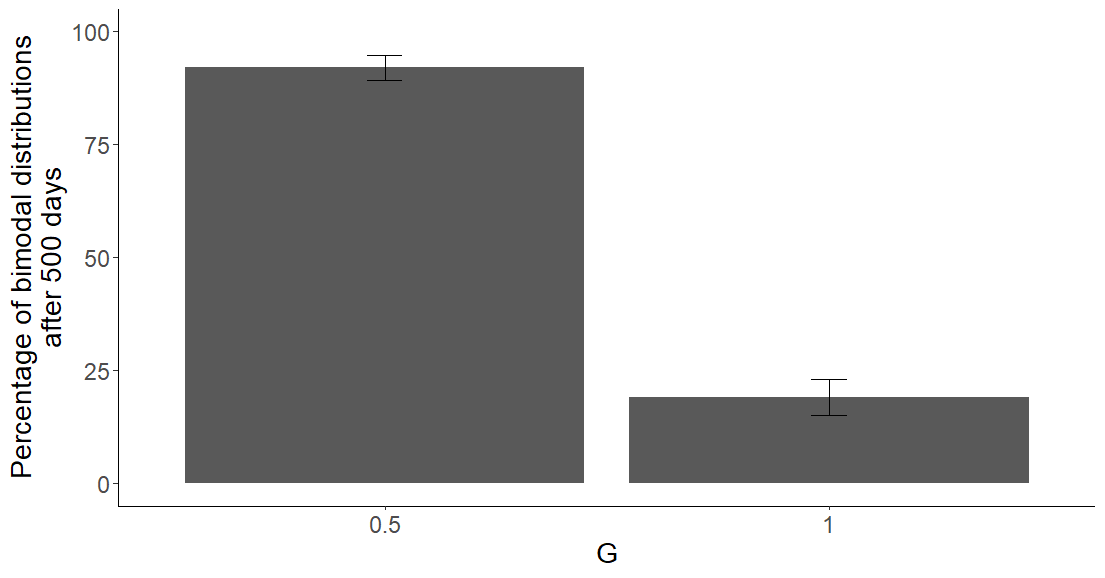

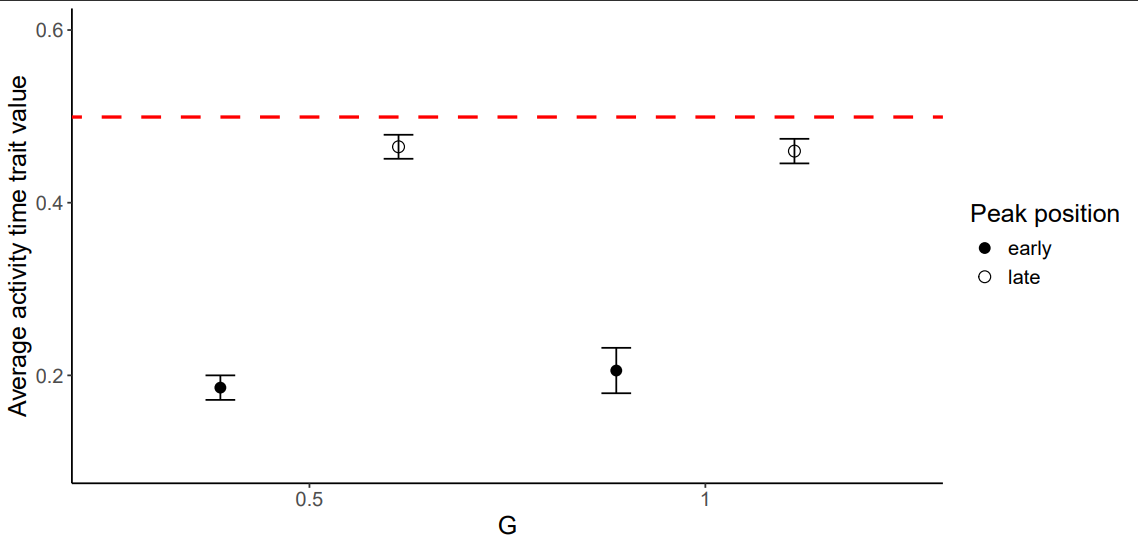

**B**

**A**

Figure S7 : **Effect of the coevolution of sexual activity timings and emergence timing on the evolution of differentiated sub-populations.** Similarly to activity timing loci, we implemented *B* = 50 emergence loci, where alleles interacted additively and had equal phenotypic effect sizes ±$\frac{1}{B}$ , chosen such that *e* values could range from 0 to 1. We measured (A) the percentage of simulations where a bimodal distribution in the timing of activities is observed at 500 days, depending on the proportion of loci *G* necessary to trigger incompatibility, and (B) the average position of early and late peaks in bimodal cases. The red dotted line indicates *t*=0.5, showing that both sub-populations displayed activity timings earlier than their ancestral state of *e* = 0.5.

Error bars are one SD. All simulations were run assuming the same parameter values: δc = 0.1, β = 3 , p = 1, K=1000.
